# Developing and Evaluating Mappings of ICD-10 and ICD-10-CM Codes to PheCodes

**DOI:** 10.1101/462077

**Authors:** Patrick Wu, Aliya Gifford, Xiangrui Meng, Xue Li, Harry Campbell, Tim Varley, Juan Zhao, Robert Carroll, Lisa Bastarache, Joshua C Denny, Evropi Theodoratou, Wei-Qi Wei

## Abstract

**Background:** The PheCode system was built upon the International Classification of Diseases, Ninth Revision, Clinical Modification (ICD-9-CM) for phenome-wide association studies (PheWAS) in the electronic health record (EHR).

**Objective:** Here, we present our work on the development and evaluation of maps from ICD-10 and ICD-10-CM codes to PheCodes.

**Methods:** We mapped ICD-10 and ICD-10-CM codes to PheCodes using a number of methods and resources, such as concept relationships and explicit mappings from the Unified Medical Language System (UMLS), Observational Health Data Sciences and Informatics (OHDSI), Systematized Nomenclature of Medicine - Clinical Terms (SNOMED CT), and National Library of Medicine (NLM). We assessed the coverage of the maps in two databases: Vanderbilt University Medical Center (VUMC) using ICD-10-CM and the UK Biobank (UKBB) using ICD-10. We assessed the fidelity of the ICD-10-CM map in comparison to the gold-standard ICD-9-CM→PheCode map by investigating phenotype reproducibility and conducting a PheWAS.

**Results:** We mapped >75% of ICD-10-CM and ICD-10 codes to PheCodes. Of the unique codes observed in the VUMC (ICD-10-CM) and UKBB (ICD-10) cohorts, >90% were mapped to PheCodes. We observed 70-75% reproducibility for chronic diseases and <10% for an acute disease. A PheWAS with a lipoprotein(a) (LPA) genetic variant, rs10455872, using the ICD-9-CM and ICD-10-CM maps replicated two genotype-phenotype associations with similar effect sizes: coronary atherosclerosis (ICD-9-CM: *P* < .001, OR = 1.60 vs. ICD-10-CM: *P* < .001, OR = 1.60) and with chronic ischemic heart disease (ICD-9-CM: *P* < .001, OR = 1.5 vs. ICD-10-CM: *P* < .001, OR = 1.47).

**Conclusions:** This study introduces the initial “beta” versions of ICD-10 and ICD-10-CM to PheCode maps that will enable researchers to leverage accumulated ICD-10 and ICD-10-CM data for high-throughput PheWAS in the EHR.

## Introduction

### Background

Electronic health records (EHRs) have become a powerful resource for biomedical research in the last decade, and many studies based on EHR data have used International Classification of Diseases (ICD) codes.[1] Often, these ICD codes are grouped to reflect biologically meaningful phenotypes and diseases.[2] Tools that group ICD codes include the Agency for Healthcare Research and Quality (AHRQ) Clinical Classification Software (CSS) and the PheCode system developed to facilitate phenome-wide association studies (PheWAS) in the EHR.[3,4] In one of the first studies to conduct a genotype-phenotype association study in the EHR, we introduced PheCodes with 733 unique codes. Subsequent studies have used PheCodes to replicate hundreds of known genotype-phenotype associations and discover dozens of new ones, some that have been validated in follow-up investigations.[5–16]

### Prior Work

We created the initial version of PheCodes on ICD Ninth Revision, Clinical Modification (ICD-9-CM) codes, thus the move to ICD-10-based codes necessitated a new PheCode map, as the ICD-10 system differs from ICD-9-CM in terms of number of codes, granularity, semantics, and organization.[17,18] In 1979, the World Health Organization (WHO) developed ICD-9 to track mortality and morbidity. To improve its application to clinical billing, the U.S. National Center for Health Statistics (NCHS) modified ICD-9 codes to create ICD-9-CM, whose end-of-life date was scheduled around the year 2000, but was delayed until October 2015.[17] In 1990, the WHO developed ICD-10,[19] which the NCHS used to create ICD-10-CM to replace ICD-9-CM. Whereas countries outside of the U.S. have used ICD-10 for many years, the U.S. moved to ICD-10-CM in October 2015. Though it is based on ICD-10, ICD-10-CM contains many more codes with increased granularity. For example, the 2018AA release of the Unified Medical Language System (UMLS)[20] contains 94,201 unique ICD-10-CM and 12,027 unique ICD-10 codes after exclusion of range codes (e.g. ICD-10-CM “A00-A09”).There are ICD-10 codes that do not exist in ICD-10-CM, and vice versa, such as ICD-10 A16.9 (Respiratory tuberculosis unspecified, without mention of bacteriological or histological confirmation), which has no ICD-10-CM equivalent. Moving from ICD-9-CM to ICD-10-CM resulted in a couple of major changes. First, the structure moved from a broadly numeric-based system in ICD-9-CM to an alphanumeric system in ICD-10-CM. Second, ICD-10-CM contains much more granular information than ICD-9-CM, such as observed with the ~10× increase in number of diabetes-related codes in ICD-10-CM.[17]

To develop the original PheCode system, we combined one or more related ICD-9-CM codes into distinct diseases or traits. For example, three depression-related ICD-9-CM codes 311, 296.31, and 296.21 are condensed to PheCode 296.2 (Depression). With the help of clinical experts in disparate domains, such as cardiology and oncology, we have updated the PheCode groupings.[21,22] An important feature of PheCodes is that it allows the identification of individuals with a phenotype of interest and the specification of accurate control populations by excluding individuals who have similar or potentially overlapping diseases (e.g. patients with codes representing type 1 diabetes or secondary diabetes mellitus would not serve as controls for type 2 diabetes).[22] PheCodes also better align with diseases mentioned in clinical practice and for genomic studies than ICD-9-CM codes and the AHRQ CCS for ICD-9.[22] The most recent iteration of PheCodes consists of 1,866 hierarchical phenotype codes that map to 15,558 ICD-9-CM codes.[23,24]

Though the PheCode system is effective at replicating and identifying novel genotype-phenotype associations, PheWAS have largely been limited to using ICD-9-CM codes. A few studies have mapped ICD-10 codes to PheCodes by converting ICD-10 to ICD-9, and then mapping the converted ICD-9 codes to PheCodes.[7,14] However, these studies limited their mappings to ICD-10 codes, did not provide an ICD-10-CM→PheCode map, and did not evaluate the accuracy of these maps.

### Goal of This Study

In this study, we developed and evaluated maps of ICD-10 and ICD-10-CM codes to PheCodes. To our knowledge, this is one of the first maps that translates ICD-10-CM to PheCodes. The primary aims of this study were to create an initial “beta” map to perform PheWAS using ICD-10 and ICD-10-CM codes and to focus the analyses on PheWAS-relevant codes. Our goal was to demonstrate that researchers should expect similar results from the ICD-10-CM→PheCode map compared to the gold-standard ICD-9-CM map. To accomplish this goal, we investigated PheCode prevalence, phenotype reproducibility, and the results from a PheWAS.

## Methods

### Databases

In this study, we used data obtained from the VUMC and UKBB[25] databases. The VUMC EHR contains records of >2.5 million unique individuals. The UKBB is a prospective longitudinal cohort study designed to investigate the genetic and environmental determinants of diseases in UK adults.[25] Between 2006-2010, the study recruited >500,000 men and women aged 40-69 years. Participants consented to allow their data to be linked to their medical records. EHR records of UKBB were obtained under an approved data request application (ID:10775).

VUMC had >2.5 years of ICD-10-CM data (~2015-10-01 to 2017-06-01), while the UKBB had >2 decades of ICD-10 data[25] (~1995-04-01 to 2015-03-31). VUMC includes codes for inpatient and outpatient encounters, whereas UKBB codes in this study are inpatient codes.

### Mapping ICD-10-CM and ICD-10 codes to PheCodes

To maximize the coverage of ICD-10-CM by PheCodes, we used the ICD-10-CM codes from the UMLS 2018AA,[20] and a number of automated methods to translate ICD-10-CM to PheCodes. An ICD-10-CM code was mapped either directly and/or indirectly to a PheCode (Figure 1), with some mappings that overlapped between the two methods. We mapped directly 515 ICD-10-CM codes to PheCodes by matching the two descriptions regardless of capitalization, e.g. ICD-10-CM H52.4 (Presbyopia)→PheCode 367.4 (presbyopia). Indirect mapping (82,287 ICD-10-CM codes) used the existing ICD-9-CM→PheCode map. To convert ICD-10-CM codes indirectly, we started with the ICD-10-CM→ICD-9-CM General Equivalence Mapping (GEM) maps.[26] Since GEM did not include all of the ICD-10-CM codes,[18] we complemented this approach with UMLS semantic mapping,[27] OHDSI concept relationships,[28,29] and NLM maps.[30]

**Figure 1.**
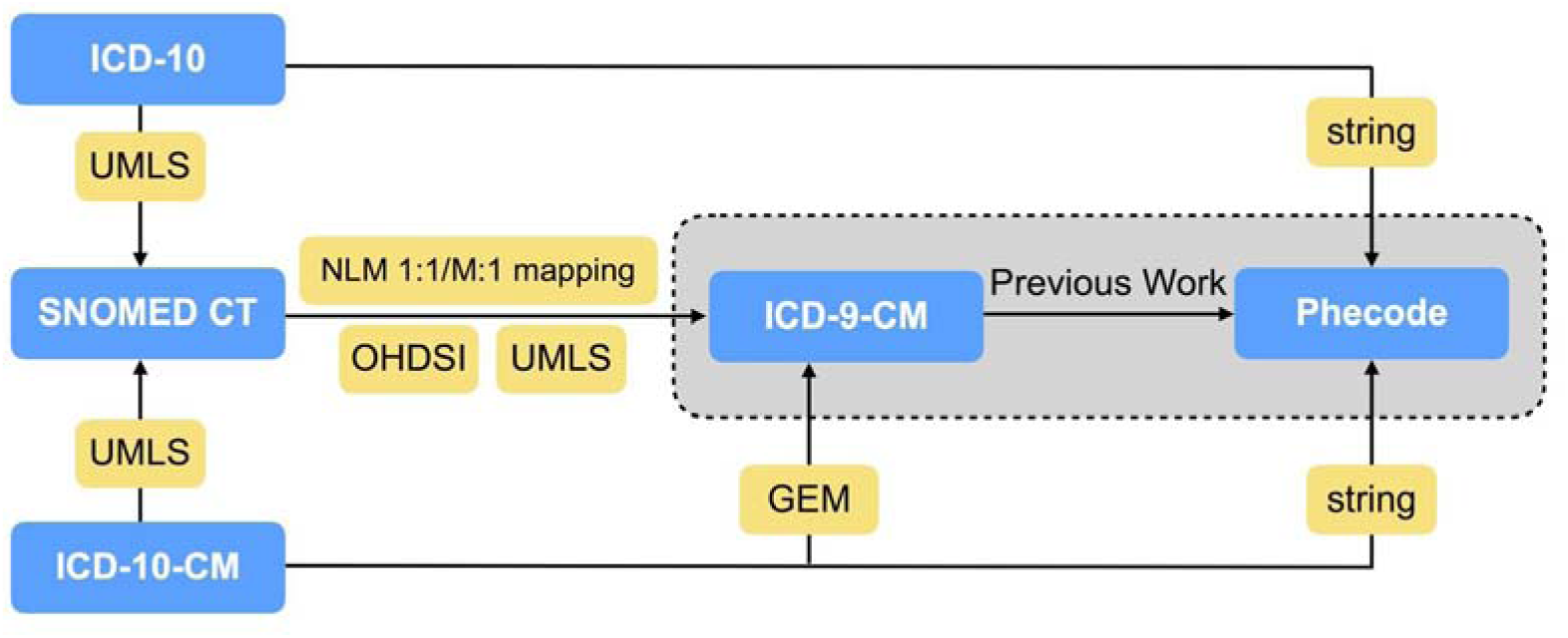
Mapping strategy for ICD-10 and ICD-10-CM codes to PheCodes. Mapping methods between code systems. The gray dashed-outline box indicates previously published work.[3,21,23] Acronyms used: SNOMED CT: Systematized Nomenclature of Medicine Clinical Terms. UMLS: Unified Medical Language System. GEM: General Equivalence Mappings. OHDSI: Observational Health Data Sciences and Informatics. NLM: National Library of Medicine. M:1: many to one.

More specifically, with GEM, we mapped ICD-10-CM→ICD-9-CM and included all mappings where the “approximate” flag was 0 or 1. The mapping of ICD-10-CM E11.9 (Type 2 diabetes mellitus without complications)→PheCode 250.2 (Type 2 diabetes) illustrates this indirect approach: ICD-10-CM E11.9→ICD-9-CM 250.0 (Diabetes mellitus without mention of complication)→PheCode 250.2. In the second approach to indirect mapping, ICD-10-CM codes were first mapped to Systematized Nomenclature of Medicine Clinical Terms (SNOMED CT) through UMLS Concept (CUI) equivalents, which were then converted to ICD-9-CM through UMLS CUI equivalents,[20,27] Observational Health Data Sciences and Informatics (OHDSI),[28] or National Library of Medicine (NLM) maps.[30] For example, ICD-10-CM L01.00 (Impetigo, unspecified)→CUI C0021099→SNOMED CT 48277006→OHDSI Concept ID 140480→OHDSI Concept ID 44832600→ICD-9-CM 684→PheCode 686.2 (Impetigo).

If an ICD-10-CM code (e.g. ICD-10-CM I10 [Essential hypertension]) was mapped to a child PheCode (e.g. PheCode 401.1 [Essential hypertension]) and its parent code (e.g. PheCode 401 [Hypertension]), we only kept the mapping to the child PheCode (i.e. PheCode 401.1) to maintain the granular meanings of the ICD-10-CM and because the PheCodes themselves are arranged hierarchically. We kept all mappings if an ICD-10-CM (e.g. ICD-10-CM D57.812 [Other sickle-cell disorders with splenic sequestration]) was mapped to multiple distinct PheCodes (e.g. PheCodes 282.5 [Sickle cell anemia] and 289.5 [Diseases of spleen]). This latter association created a polyhierarchical nature to PheCodes that did not exist previously. The ICD-10-CM→PheCode map is hosted at https://phewascatalog.org/phecodes_icd10cm.[31]

We used ICD-10 from the UMLS 2018AA.[20] ICD-10 codes were mapped to PheCodes in a similar manner to ICD-10-CM, except that a ICD-10→ICD-9-CM GEM did not exist, so we used only a combination of string matching, UMLS, NLM, OHDSI, and SNOMED CT resources (Figure 1). The ICD-10→PheCode map is hosted at https://phewascatalog.org/phecodes_icd10.[32]

### Evaluation of PheCode coverage of ICD-10 and ICD-10-CM in UKBB and VUMC

To evaluate the PheCode coverage of ICD-10 and ICD-10-CM in UKBB and VUMC, respectively, we calculated the number of codes in the two official maps, the number of codes mapped to PheCodes, and the number of mapped and unmapped official codes used in the two EHRs (Figure 2).

**Figure 2.**
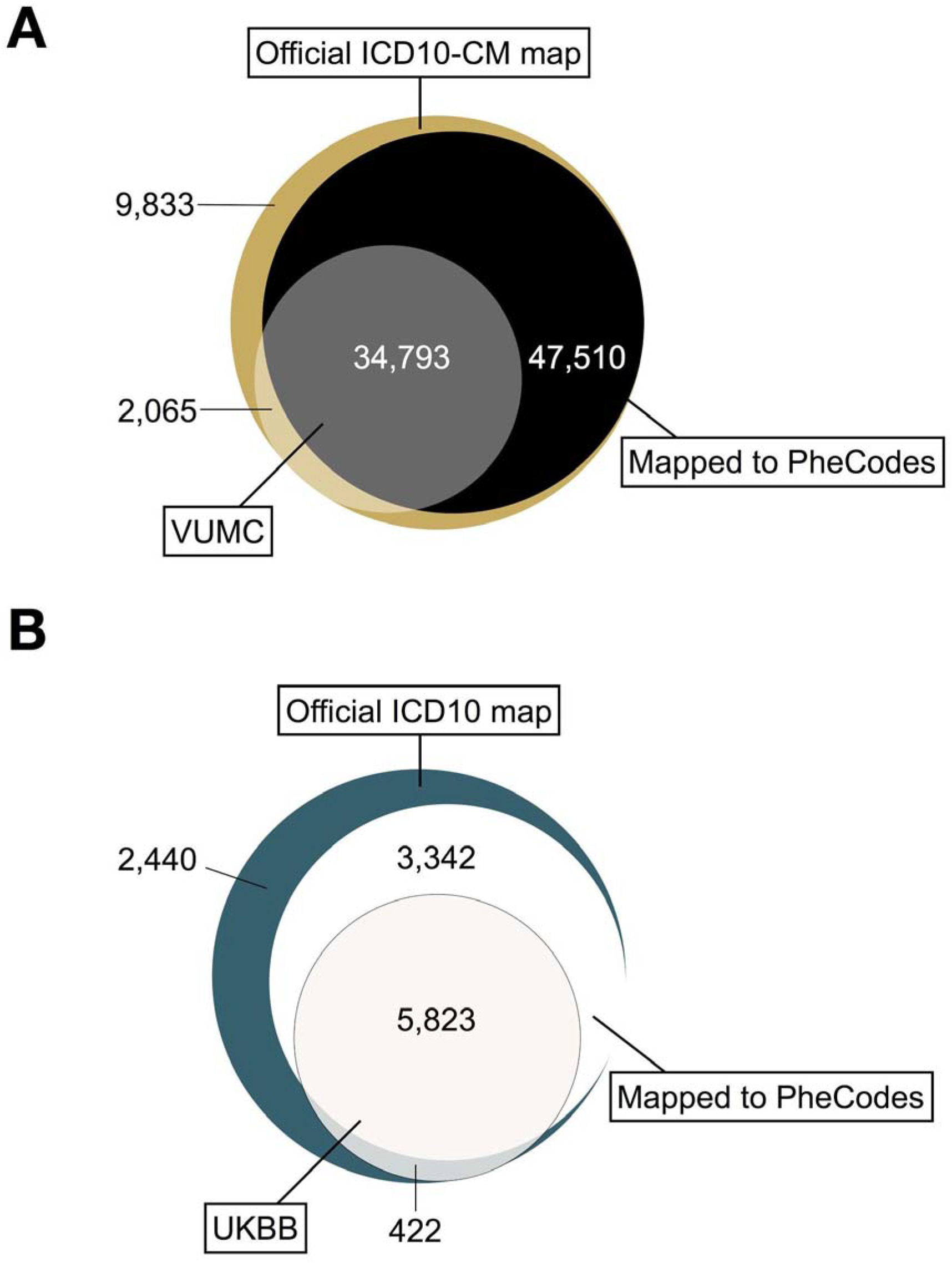
Counts of distinct ICD-10-CM codes at VUMC and ICD-10 codes in UKBB. (A) Number of unique ICD-10-CM codes in each category. For example, there are 34,793 unique codes (grey section) that are in the official ICD-10-CM system, observed in the Vanderbilt University Medical Center (VUMC) dataset, and mapped to PheCodes. (B) Number of unique ICD-10 codes in each category. For example, there are 5,823 unique codes (off-white section) that are in the official ICD-10 system, observed in the UK Biobank (UKBB) dataset, and mapped to PheCodes.

To identify potential limitations of our automated mapping method, authors with a clinical background (P.W., W.Q.W.) manually reviewed the unmapped ICD-10 and ICD-10-CM codes that were used at UKBB and VUMC, respectively.

### Reproducibility analysis of ICD-10-CM→PheCode map

We aimed to provide evidence that the ICD-10-CM→PheCode map results in phenotypes similar to ones sourced from the ICD-9-CM→PheCode map. First, we selected 357,728 patients in the VUMC EHR who have ≥1 ICD-9-CM and ≥1 ICD-10-CM codes in two 18-month windows. We selected windows to occur prior to and after VUMC’s transition to ICD-10-CM. To reduce potential confounders, we left a six-month buffer after ICD-9-CM was replaced with ICD-10-CM. Further, the ICD-10-CM observation window ended before VUMC switched from its locally developed EHR[33] to the Epic system. This created two windows ranging from 2014-01-01→2015-06-30 for ICD-9-CM, and 2016-01-01→2017-06-30 for ICD-10-CM (Figure 3). The final cohort consisted of 55.1% female with mean (standard deviation, SD) 45 (25) years old. We extracted all ICD-9-CM and ICD-10-CM codes from these two windows for each patient. We mapped these codes to PheCodes using the ICD-9-CM→PheCode map [23] and the ICD-10-CM→PheCode map.

**Figure 3.**
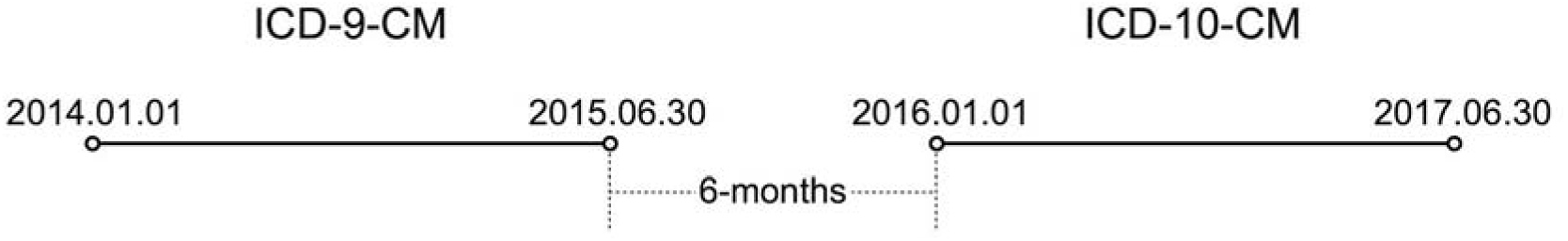
Timeline of the two 18-month periods from which the ICD-9-CM and ICD-10-CM codes were analyzed at VUMC. The cohort of 357,728 patients had at least one ICD-9-CM and one ICD-10-CM code in the respective 18-month windows.

We hypothesized that in comparison to the gold-standard ICD-9-CM→PheCode map, the new ICD-10-CM→PheCode map created a reproducible set of phenotype definitions. To test this hypothesis, we designed an experiment to see whether PheCodes had assigned consistently the same set of phenotypes representing chronic diseases to patients in the ICD-10-CM period as when ICD-9-CM was in use. We chose four chronic diseases (Hypertension, Hyperlipidemia, Type 1 Diabetes, and Type 2 Diabetes) and as a negative control, one acute disease (Intestinal infection). We expected high phenotype reproducibility for the chronic diseases and low phenotype reproducibility for the acute disease. PheCodes 401.* (“*” means one or more digits or a period) represented Hypertension, 272.* represented Hyperlipidemia, 250.1* represented Type 1 diabetes, 250.2* represented Type 2 diabetes, and 008.* represented Intestinal infection.

For each phenotype, we reported the number of cases in the ICD-9-CM era and the number of those individuals who were also cases in ICD-10-CM. To investigate the possible reasons for individuals who did not “reproduce” a PheCode, authors with a clinical background (P.W., W.Q.W.) manually reviewed medical records of ten randomly selected patients who did not have the PheCode of interest in the ICD-10-CM era from the ‘Hypertension’, ‘Hyperlipidemia’, and ‘Type 2 diabetes’ groups, for a total of thirty patients.

### Comparative PheWAS analysis of lipoprotein(a) (LPA) single-nucleotide polymorphism (SNP)

To evaluate the accuracy of the ICD-10-CM→PheCode map, we performed two PheWAS on a LPA genetic variant (rs10455872) using mapped PheCodes from ICD-9-CM and ICD-10-CM. We recently demonstrated that rs10455872 is associated with increased risks of developing hyperlipidemia and cardiovascular diseases.[34,35]

We used data from BioVU, the de-identified DNA biobank at VUMC to conduct the PheWAS.[36] We identified 13,900 adults (56.9 % female, 59 (15) years old in 2014), who had rs10455872 genotyped, and at least one ICD-9-CM and ICD-10-CM code in their respective time windows. We observed 86.7% AA, 12.8% AG, and 0.5% GG for rs10455872. We used 1,632 PheCodes that overlapped in the time windows for PheWAS using the R PheWAS package[24] with standard logistic regression, adjusting for age, sex, and race.

## Results

### PheCode coverage of ICD-10-CM and ICD-10 in VUMC and UKBB

Of the official ICD-10-CM codes,[20] 82,303 (87.37%) mapped to at least one PheCode, with 7,881 (8.37%) mapping to >1 PheCode. Of all possible ICD-10 codes, 9,165 (76.20%) mapped to at least one PheCode, and 289 (2.40%) mapped to >1 PheCode. For example, ICD-10-CM E13.36 (Other specified diabetes mellitus with diabetic cataract) mapped to three PheCodes: 366 (Cataract), 250.7 (Diabetic retinopathy), and 250.23 (Type 2 diabetes with ophthalmic manifestations). Whereas, ICD-10 code B21.1 (HIV disease resulting in Burkitt’s lymphoma) maps to two PheCodes: 071.1 (HIV infection, symptomatic) and 202.2 (Non-Hodgkin’s lymphoma).

Among the 36,858 ICD-10-CM codes used at VUMC, 34,793 (94.40%) codes were mapped to PheCodes. In the UKBB, 5,823 (93.24%) of the ICD-10 codes mapped to PheCodes (Table 1, Figure 2). Considering all the instances of ICD-10-CM and ICD-10 codes used at each site, we generated a total count of unique codes grouped by patient and date, and a subset of the codes that mapped to PheCodes (Table 1). Among the total number of codes used, 89.72% of ICD-10-CM and 83.68% of ICD-10 codes were mapped to PheCodes.

**Table 1.** ICD-10-CM and ICD-10 codes data summary.

|  | ICD-10-CM<sup>a</sup> (No.) | ICD-10<sup>b</sup> (No.) |
| --- | --- | --- |
| **Official classification systems** |  |  |
| Unique codes | 94,201 | 12,027 |
| **Unique codes mapped** | 82,303 (87.37%) | 9,165 (76.20%) |
| **Official codes used in cohorts** |  |  |
| **Unique codes** | 36,858 | 6,245 |
| **Unique codes mapped** | 34,793 (94.40%) | 5,823 (93.24%) |
| **Total patients (with ICD-10-CM or ICD-10 codes)** | 651,649 | 391,181 |
| **Total instances of all ICD codes** | 19,682,697 | 5,114,363 |
| **Instances mapped to PheCodes** | 17,658,470 (89.72%) | 4,279,544 (83.68%) |

<sup>a</sup>ICD-10-CM data for patients derived from Vanderbilt University Medical Center (VUMC) included inpatient and outpatient data.
<sup>b</sup>ICD-10 inpatient data derived from UK Biobank.

### Analysis of unmapped ICD-10 and ICD-10-CM codes

Majority of the unmapped ICD-10 codes used in the UKBB dataset represented medical concepts related to personal (i.e. past medical history) or family history of disease. For ICD-10-CM, removing codes used at VUMC that we expected to be unmapped (i.e. local or supplementary classification codes) left 2,065 ICD-10-CM codes that did not map to a PheCode. Six hundred seventy codes remained after excluding X, Y, and Z codes (1,395 codes), majority of which represented either “external causes of morbidity” or “factors influencing health status and contact with health services”. All of the remaining unmapped ICD-10-CM codes in this cohort had <200 unique individuals (i.e. <.1% of the cohort), and majority of the ICD-10-CM codes with >10 unique individuals do not represent biologically or clinically meaningful phenotypes relevant to genotype-phenotype association studies. For example, 287 (59.2%) of the unmapped ICD-10-CM codes represented external causes of morbidity, such as assault and injuries due to motor vehicle accidents.

### Reproducibility analysis of the ICD-10-CM→PheCode map

In the defined 18-month time windows, a cohort 357,728 patients had both ICD-9-CM and ICD-10-CM codes (Figure 3). For the chronic diseases, 70-75% of individuals with the relevant PheCodes in the ICD-9-CM observation period also had the same PheCodes of interest during the ICD-10-CM period. On the contrary, for the reproducibility analysis with an acute disease, we observed that <10% of individuals who had PheCodes 008.* (Intestinal infection) in the ICD-9-CM period also had the same PheCodes in the ICD-10-CM period (Table 2).

**Table 2.** ICD-10-CM PheCode map reproducibility analysis.

| Phenotype | PheCodes<sup>a</sup> | #Cases in ICD-9-CM | Number of Individuals (%),<br>(Case in ICD-10-CM \| Case in ICD-9-CM)<sup>b</sup> |
| --- | --- | --- | --- |
| Hypertension | 401.* | 65,216 | 49468 (75.85%) |
| Hyperlipidemia | 272.* | 51,187 | 36187 (70.70%) |
| Type 1 diabetes | 250.1* | 5,782 | 4,412 (76.31%) |
| Type 2 diabetes | 250.2* | 25,077 | 19,066 (76.03%) |
| Intestinal infection | 008.* | 3,410 | 273 (8.01%) |

<sup>a</sup>In the PheCode column, “\*” means one or more digits or a period, e.g. PheCode 401.\* = PheCode [401, 401.1, 401.3, 401.22, 401.21, 401.2].
<sup>b</sup>In the last column, “Case in ICD-10-CM | Case in ICD-9-CM” indicates patients who had phenotype of interest in ICD-9-CM period also had the same phenotype during the ICD-10-CM period.

To investigate the possible reasons for individuals who did not “reproduce” a PheCode, we manually reviewed medical records of ten randomly selected patients who did not have the PheCode of interest in the ICD-10-CM period from the ‘Hypertension’, ‘Hyperlipidemia’, and ‘Type 2 diabetes’ groups. In general, these patients did not “reproduce” their phenotypes in the ICD-10-CM era, because they were not given ICD-10-CM codes corresponding with the phenotypes. Reasons for not being given these codes include a short ICD-10-CM observation window of 18 months, false-positive phenotyping in the ICD-9-CM era (e.g. patient with Type 1 diabetes given Type 2 diabetes ICD-9-CM code), visits limited to specialists (e.g. only seen by neurologist in ICD-10-CM period), and limited number of visits (<2) to VUMC during ICD-10-CM period.

### Comparative PheWAS analysis of LPA SNP, rs10455872

To further evaluate the ICD-10-CM→PheCode map, we performed and compared the results of a PheWAS for rs10455872. One PheWAS was conducted using the ICD-9-CM map and another was conducted using the ICD-10-CM map. Both analyses replicated previous findings, showing statistically significant (alpha=.05) positive correlations between the minor allele of rs10455872 and coronary atherosclerosis (ICD-9-CM: *P* < .001, OR = 1.60 vs. ICD-10-CM: *P* < .001, OR = 1.60) and with chronic ischemic heart disease (ICD-9-CM: *P* < .001, OR = 1.56 vs. ICD-10-CM: *P* < .001, OR = 1.47) (Figure 4).

**Figure 4.**
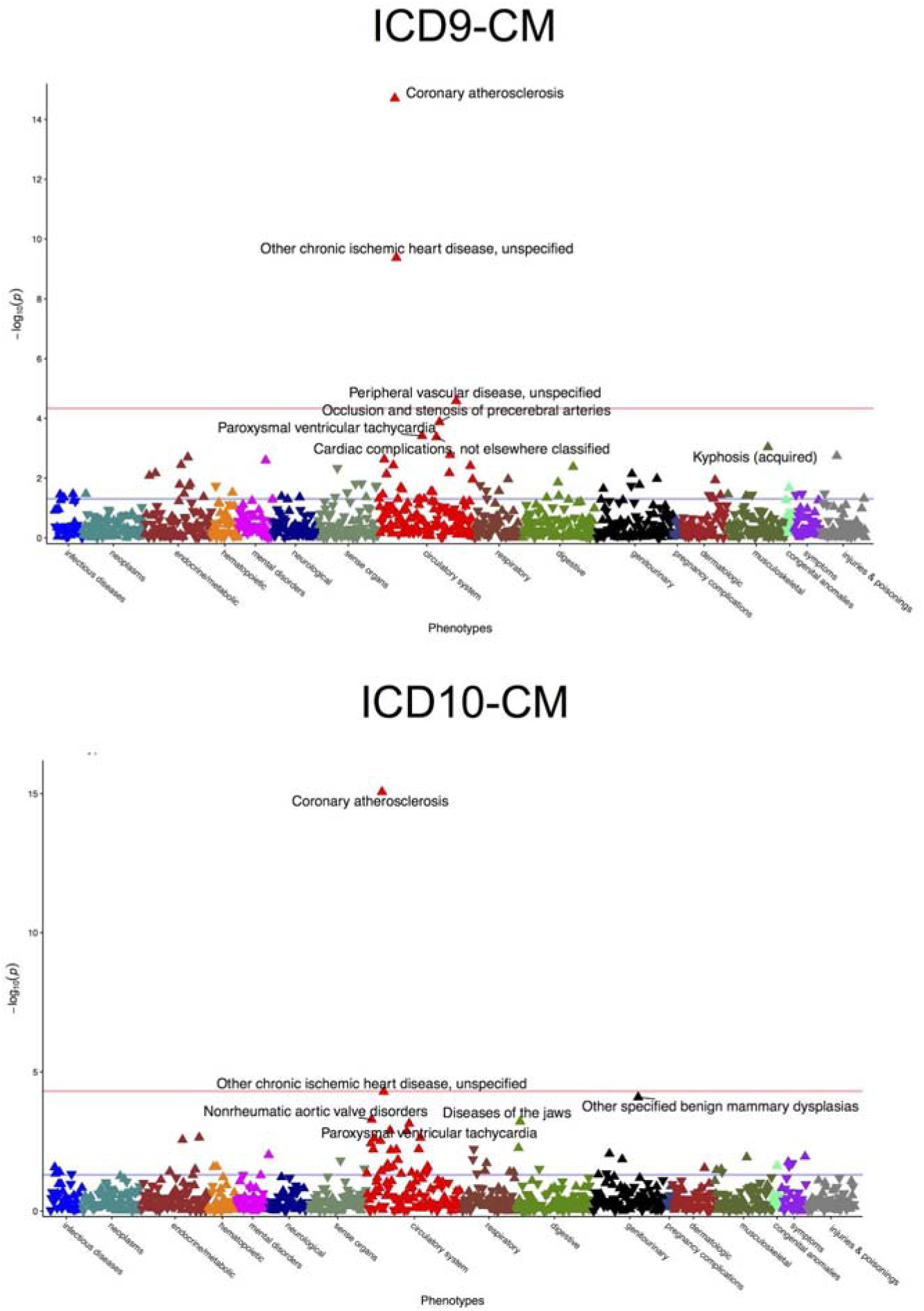
Comparative PheWAS of lipoprotein(a) (LPA) genetic variant, rs10455872. Coronary atherosclerosis (PheCode 411.4) and other chronic ischemic heart disease (PheCode 411.8) were top hits associated with rs10455872 in a phenome-wide association study (PheWAS) analysis conducted using ICD-9-CM (top) and ICD-10-CM (bottom) to PheCode maps. Analysis was adjusted for age, sex, and race.

## Discussion

### Main findings: Maps of ICD-10 and ICD-10-CM codes to PheCodes have high coverage and yield similar results as the ICD-9-CM→PheCode map

This work investigated the results of mapping ICD-10 and ICD-10-CM codes to PheCodes and evaluation in two databases. These results show that the majority of billed instances can be mapped to PheCodes using ICD-10 or ICD-10-CM codes. Our analyses suggest that researchers can expect that phenotypes sourced using the ICD-10-CM→PheCode map to be similar to those sourced from the ICD-9-CM→PheCode map. As the use of ICD-10 and ICD-10-CM codes increases, so does the need for convenient and reliable methods of aggregating codes to represent biologically meaningful phenotypes for biomedical research. Since the introduction of PheCodes, many studies have demonstrated the value of aggregating ICD-9-CM codes. This resource will allow biomedical researchers to leverage clinical data represented by ICD-10 and ICD-10-CM codes for their large-scale genotype-phenotype association studies in the EHR.

### ICD-10 and ICD-10-CM codes not mapped to PheCodes

Analysis of the unmapped ICD-10 codes demonstrates a possible area of expansion for PheCodes. The ICD-10→PheCode map missed many medical concepts representing personal or family history of disease.

We observed that a majority of the unmapped ICD-10-CM codes represented concepts that we did not expect to have PheCode equivalents. Majority of the codes were from ICD-10-CM chapters 20 (External causes of morbidity) and 21 (Factors influencing health status and contact with health services). Codes from chapter 19 (Injury, poisoning, and certain other consequences of external causes) also made up a large fraction of unmapped codes, such as ICD-10-CM T38.3×6A (Underdosing of insulin and oral hypoglycemic [antidiabetic] drugs, initial encounter). We did not expect ICD-10-CM T38.3×6A to map to a PheCode, as it is an “encounter” code that is not relevant to PheWAS. Three-digit codes that are not frequently used for billing purposes, such as ICD-10-CM I67 (Other cerebrovascular diseases), also made up a large number of unmapped codes. A few potentially meaningful phenotypes, such as ICD-10-CM O04.6 (Delayed or excessive hemorrhage following [induced] termination of pregnancy), are also unmapped and represent areas of potential expansion for PheCodes.

### ICD-10-CM→PheCode map phenotype reproducibility analysis

In general, our analysis suggests that majority of the PheCodes that are not reproduced in the ICD-10-CM observation period are not due to inaccurate mappings in the ICD-10-CM→PheCode map. This study’s reproducibility analysis (Table 2) demonstrates that the vast majority of patients (70-75%) with PheCodes of four chronic diseases sourced from ICD-9-CM codes will also have the same phenotypes in the ICD-10-CM era. In comparison, when the same experiment is repeated for an acute disease (Intestinal infection), a small minority (<10%) of patients will have the same phenotype in the ICD-10-CM period.

Using the ICD-9-CM and ICD-10-CM maps, PheWAS found similar statistically significant associations and effect sizes with coronary atherosclerosis and chronic ischemic heart disease (Figure 4). Results of this analysis provide additional support for the accuracy of the ICD-10-CM map when compared to the gold-standard ICD-9-CM→PheCode map.

### PheWAS using ICD-10→PheCode map

Two published studies have tested the ICD-10→PheCode map to identify genotype-phenotype associations using data from the UKBB. Zhou et al. used the map to demonstrate a method that adjusts for case-control imbalances in a large genome-wide PheWAS.[37] Li et al. used the same map to estimate the causal effects of elevated serum uric acid across the phenome.[16]

### Related work

The research community has developed alternative models to create patient cohorts in the EHR by collapsing ICD codes into clinically meaningful phenotypes.[33,34] For example, OHDSI employs the OMOP Common Data Model (CDM) to represent data consistently across disparate systems and supports researchers interested in using observational health data.[35] In contrast, PheCodes were designed for high-throughput PheWAS in the EHR and has the inherent ability to define precise controls using built-in exclusion criteria.[22] For example, researchers who want to use PheCode 250.2 for Type 2 diabetes can automatically exclude individuals with Type 1 diabetes using PheCode 250.1 from the control population. The new PheCode maps maintain this feature to identify high-quality control cohorts. Since we used OHDSI concept relationships to create the new PheCode maps, there may be overlap of individuals between PheCode-defined cohorts and those created with source data in ICD-10 or ICD-10-CM codes using the OMOP CDM. While researchers can use OHDSI concept relationships for PheWAS, we believe that PheCodes are more suitable for high-throughput genotype-phenotype association studies in the EHR.

### Limitations

This study has limitations. First, due in part to the mapping strategy that we used to map ICD-10→ICD-9-CM, only 85.53% (1596/1866) of the PheCodes are mapped to an ICD-10 code. Second, the VUMC data are from a single site, thereby making it difficult to generalize the results of our accuracy studies (e.g. phenotype reproducibility analysis and LPA SNP PheWAS) to patient cohorts in other EHRs. Third, this work did not aim to thoroughly evaluate the accuracy of these “beta” PheCode maps, so our assumptions about the high accuracy of the resources used in mapping could have affected the accuracy of the new PheCode maps. In the 2009 ICD-10-CM→ICD-9-CM GEM, >90% of the mappings are “approximate”.[17] For this study’s purposes, we included all mappings where the “approximate” flag was 0 or 1 in the 2018 ICD-10-CM to ICD-9-CM GEM, which was a trade-off between coverage and mapping accuracy.

Fourth, our automated approach to map >80,000 ICD-10-CM and >9,000 ICD-10 codes to PheCodes with minimal human-engineering could have affected the accuracy of the final maps. Hripcsak et al.[38] recently evaluated the effects of translating ICD-9-CM codes to SNOMED CT codes on the creation of patient cohorts. In general, they found that mapping source billing codes to a standard clinical vocabulary (e.g. ICD-9-CM→SNOMED CT) did not greatly affect cohort selection. Their findings suggested that optimized domain knowledge-engineered mappings outperform simple automated translations between clinical vocabularies. Using four phenotype concept sets, they showed that automated mappings resulted in errors of up to 10% and domain-knowledge engineered mappings to have errors of <.05%. Other studies have also found that mapping performance is generally better with smaller value sets.[18] Future mapping studies could consider using an iterative forward and backward mapping approach using GEMs to create a more comprehensive and accurate map between ICD-9-CM and ICD-10-CM.[18]

### Future Directions

Currently, if an ICD-10 or ICD-10-CM code maps to ≥2 codes unlinked PheCodes, we keep all of the mappings. In subsequent studies, it will be important to further scrutinize these mappings to ensure accuracy through manual review. Future work involves expanding and updating the mappings to PheCodes, addressing the unmapped codes, and extensive manual validation of mapping accuracy with input from the research community.

### Conclusions

In this paper, we introduced our work on mapping ICD-10 and ICD-10-CM codes to PheCodes. We provide initial “beta” maps with high coverage of EHR data in two large databases. Results from this study suggested that the ICD-10-CM→PheCode map created phenotypes similar to those generated by the ICD-9-CM→PheCode map. These mappings will enable researchers to leverage accumulated ICD-10 and ICD-10-CM data in the EHR for large PheWAS.

## Supporting information

icd10cm_codes.csv

genotypes.csv

icd10cm_phecode_example_070319.html

## Acknowledgements

P.W., A.G., J.C.D., and W.Q.W. contributed to the design of the studies. P.W., A.G., X.M., X.L., H.C., E.T, T.V., J.Z., J.C.D., and W.Q.W. analyzed the data. P.W. and A.G. were responsible for the literature review. A.G., X.M., X.L., and E.T. retrieved the raw data. P.W., A.G., R.C., L.B., J.C.D., E.T., and W.Q.W interpreted the data. P.W., A.G., J.C.D., and W.Q.W drafted the initial manuscript. P.W., A.G., J.C.D., and W.Q.W were involved in the creation and design of figures and tables. All authors revised the document and gave final approval for publication. The project was supported by NIH grant R01 LM 010685, R01 HL133786, T32 GM007347, T15 LM007450, P50 GM115305, and AHA Scientist Development Grant 16SDG27490014. The dataset used in the analyses described were obtained from Vanderbilt University Medical Center’s BioVU, which is supported by institutional funding and by the Vanderbilt CTSA grant ULTR000445 from NCATS/NIH. This research was also conducted using the UK Biobank Resource under Application Number 10775. The work conducted in Edinburgh was supported by funding for the infrastructure and staffing of the Edinburgh CRUK Cancer Research Centre. E.T. is supported by a CRUK Career Development Fellowship (C31250/ A22804). X.M. and X.L. are supported by China Scholarship Council studentships.

## Conflicts of Interest

None declared

## Abbreviations

EHR: electronic health record
ICD: International Classification of Diseases
AHRQ: Agency for Healthcare Research and Quality
CCS: Clinical Classification Software
PheWAS: phenome-wide association studies
CM: Clinical Modification
WHO: World Health Organization
NCHS: National Center for Health Statistics
UMLS: Unified Medical Language System
GEM: General Equivalence Mapping
SNOMED CT: Systematized Nomenclature of Medicine Clinical Terms
CUI: Concept Unique Identifier
OHDSI: Observational Health Data Sciences and Informatics
CDM: Common Data Model
NLM: National Library of Medicine
VUMC: Vanderbilt University Medical Center
UKBB: UK Biobank
OR: odds ratio
LPA: lipoprotein(a)
SNP: single nucleotide polymorphism
M:1: many to one
SD: standard deviation

## Notes

#### Summary of Updates

1. Updated with example of PheWAS from ICD-10-CM codes using R PheWAS package v1.0 (https://github.com/PheWAS/PheWAS). The R code is in icd10cm_phecode_example_070319.html. Also uploaded 'icd10cm_codes.csv' and 'genotypes.csv' file used in example. 2. No change to manuscript

