## Supplementary material for "Developing and Evaluating Mappings of ICD-10 and ICD-10-CM Codes to PheCodes": icd10cm_phecode_example_070319.html

Example PheWAS with Phecode v1.2 ICD-10-CM code map beta 1


### Example PheWAS with Phecode v1.2 ICD-10-CM code map beta 1

#### Install PheWAS

We first need to install and load the v1.0 PheWAS package:

```
if(!require(PheWAS)|packageVersion("PheWAS")<'0.99'){
  if(!require(devtools)){
    install.packages("devtools")
  }
  devtools::install_github("PheWAS/PheWAS", ref='v1.0')
  library(PheWAS)
}
if(!require(readr)) {install.packages("readr")} #For faster file loading
```

#### Load data

There are a number of ways to load data, and we may have a variety of data sources for our analysis. It may be necessary to set or change your working directory (setwd). Alternatively, you can specify the complete path to the file you wish to load. Note that knitr requires us to use a special setup for the working directory above.

```
setwd("~/")
```

Next, we read our billing code data from a csv file. We specify the data types of the columns as R will want to read ICD10CM codes as numeric values.

```
library(PheWAS)
```

```
## Loading required package: dplyr
```

```
## Warning: package 'dplyr' was built under R version 3.5.2
```

```
## 
## Attaching package: 'dplyr'
```

```
## The following objects are masked from 'package:stats':
## 
##     filter, lag
```

```
## The following objects are masked from 'package:base':
## 
##     intersect, setdiff, setequal, union
```

```
## Loading required package: tidyr
```

```
## Warning: package 'tidyr' was built under R version 3.5.2
```

```
## Loading required package: ggplot2
```

```
## Warning: package 'ggplot2' was built under R version 3.5.2
```

```
## Loading required package: parallel
```

```
library(readr)
icd10cm_codes=read_csv("icd10cm_codes.csv", col_types="ifci")
```

Lastly, we will import `genotypes.csv`: contains genotype status, rsEXAMPLE for individuals in `icd10cm_codes.csv`.

```
genotypes <- read.csv('genotypes.csv',sep=',',colClasses=c("integer","integer"))
```

#### Prepare Data

Using the raw format data from plink means our genotype data will be ready to go. We are not going to transform our covariates either, so that leaves us to set up our phenotypes. The below call will use the default minimum code count of 2. This may take a few minutes to complete. Note that we are restricting some phenotypes that are specific to males or females, eg prostate cancer. We also define an aggregate function here, as we are using count data and not dates of occurrence (the preferred input).

```
phenotypes=createPhenotypes(icd10cm_codes)
```

```
## Mapping codes to phecodes...
```

```
## Aggregating codes...
```

```
## Mapping exclusions...
```

```
## Coalescing exclusions and min.code.count as applicable...
```

```
## Reshaping data...
```

#### Run PheWAS

```
results=phewas(phenotypes,genotypes,cores=1,significance.threshold=c("bonferroni"))
```

```
## Merging data using these shared columns:  id
```

```
## Finding associations...
```

```
## Compiling results...
```

```
## Cleaning up...
```

```
## Finding significance thresholds...
```

#### View Our Results

Checking out our results both as a table and figure can be helpful. There are many ways to order and view the results in R:

```
results_d=addPhecodeInfo(results)
#List the significant results
sig_results <- results_d[results_d$bonferroni&!is.na(results_d$p),]
DT::datatable(sig_results)
```

#### Plot the Results

Creating a rough visualization is straight forward.

```
phewas_plot <- phewasManhattan(results, OR.direction = T, title="My Example PheWAS Manhattan Plot", annotate.size=3)
```

```
## Scale for 'shape' is already present. Adding another scale for 'shape',
## which will replace the existing scale.
```

```
phewas_plot
```
